# Quantitative assessment of Newcastle disease virus proteins interactions with all known mucin types of Chicken and Quail

**DOI:** 10.1101/2024.01.16.575848

**Authors:** Hazem Almhanna, Arun HS Kumar

## Abstract

**Background:** Newcastle Disease (ND), caused by the Newcastle Disease Virus (NDV) poses a significant threat to poultry, leading to severe economic losses. Understanding the molecular interactions between NDV proteins and avian mucins is crucial for developing targeted interventions.

**Material and Methods:** In this study, twelve NDV proteins were systematically assessed for their interactions with sixteen quail and eight chicken mucin types, revealing diverse and species-specific binding patterns.

**Results:** High-affinity interactions between mucins (Muc5A, Muc5B, and Muc6) and NDV hemagglutinin-neuraminidase, was observed in addition to significant interactions with NDV fusion glycoprotein. Notably, chicken Muc4 displayed mid-range interactions exclusively with NDV fusion glycoprotein, highlighting potential species-specific differences in viral entry mechanisms between quails and chickens. Furthermore, the study investigated the number of binding sites on NDV proteins and chicken/quail mucins. Chicken Muc5B emerged as a standout with the highest number (20) of binding sites, suggesting its crucial role in NDV infection. The binding site analysis identified key regions in NDV fusion glycoprotein and hemagglutinin-neuraminidase, indicating potential targets for vaccine development.

**Conclusion:** This study provides a foundation for future research into optimizing diagnostic approaches and therapeutic strategies for NDV infections. Validation of these interactions with real-world clinical data, coupled with an exploration of tissue-specific mucin expression patterns, could further enhance our understanding of host-virus dynamics. The identified interactions offer promising avenues for developing vaccines that target specific binding sites, thereby contributing to the effective control and prevention of Newcastle Disease in poultry populations.

## Introduction

Newcastle disease (ND), also known as avian paramyxovirus or avian pneumoencephalitis, is a highly contagious viral infection that affects birds, particularly poultry.^1^ It is caused by the Newcastle disease virus (NDV, Avian paramyxovirus 1), which belongs to Avulavirus genus and the Paramyxoviridae family, it has a ribonucleic acid (RNA) genome.^2^ Paramyxoviruses have different basic attachment proteins including hemagglutinin-neuraminidase (HN), hemagglutinin (HA), glycoprotein (G), and fusion (F) proteins which might bind receptors with host cell plasma membranes.^3,4^ The disease can affect various species of birds, including chickens, turkeys, ducks, pigeons, and wild birds^5^, and can be transmitted through direct contact with infected birds, contaminated surfaces, or through the respiratory route, such as through droplets or aerosols.^5^

Symptoms of ND can vary depending on the severity of the infection, but they commonly include respiratory signs like coughing, sneezing, nasal discharge, and difficulty breathing. Other symptoms may include decreased egg production, diarrhoea, nervous signs like tremors or paralysis, and even sudden death.^6,7^ ND can manifest in various forms, ranging from mild to severe and the severity of the disease depends on the strain of the NDV and the susceptibility of the bird species affected.^8^ ND displays different forms including asymptomatic or subclinical form, and birds infected would show no visible symptoms or only very mild signs.^8^ Also, respiratory form is characterized by respiratory symptoms such as coughing, sneezing, nasal discharge, and difficulty breathing, in addition to conjunctivitis or swelling around the eyes.^5,8^ The digestive form is more common with young birds in which infected birds would suffer from diarrhoea, greenish faeces, and reduced feed intake.^2,5,8^ While the nervous or neurological form is characterized by symptoms such as tremors, paralysis, twisted necks, circling behaviour, or even complete loss of motor control.^2,5,8^ However the viscerotropic form is the most severe and deadly form of ND, and causes high mortality rates.^5,8^ In this form infected birds may exhibit a combination of respiratory, digestive, and neurological symptoms.^5,8,9^

NDV primarily affects birds by interacting with specific receptors on the surface of host cells. The main receptor involved in the entry of NDV into cells is sialic acid. NDV recognizes and binds to sialic acid receptors on the host cell surface through a protein on its own viral envelope called the hemagglutinin-neuraminidase (HN) protein.^8-10^ Different strains of NDV may have preferences for specific types of sialic acid receptors, some strains preferentially bind to sialic acid receptors with an alpha-2,3 linkage, while others have a preference for alpha-2,6-linked sialic acids.^8-10^ Mucins are large, heavily glycosylated proteins that play a critical role in maintaining the protective mucus layer in various tissues, including the respiratory tract, gastrointestinal tract, and reproductive system.^11-14^ Mucins are divided into a gel-forming mucin that is responsible for the production of mucus in the respiratory and digestive tracts such as MUC5A, and MUC5B, and a membrane-bound mucin is expressed on the cell surfaces such as MUC4, and MUC6.^15,16^ Mucins are glycoproteins connected to different types of sialic acids.^17-19^ Therefore, mucins are involved in maintaining the integrity of the respiratory tract and protecting it from pathogens, irritants, and dehydration.^19,20^ MUC4 acts as a physical barrier and helps prevent the attachment and invasion of microorganisms.^10,19-21^ As sialic acids are structurally integral to mucins playing important roles in various biological processes, such as cell adhesion, signalling, and protection of the mucosal surfaces, we hypothesised that surface proteins of NDV will have interactions with mucins. To assess this hypothesis, this study objectively assessed the interactions of twelve different NDV proteins with all known mucin types in quails and chickens.

## Material and Methods

### Newcastle disease virus proteins

The UniProt database was reviewed with the following key words; Newcastle disease, Avian orthoavulavirus 1, quail and chicken, to identify all available proteins of Newcastle disease virus. After systematic assessment to identify duplicates and close isoforms, we shortlisted twelve NDV proteins for further assessment in this study. These proteins are: 1. RNA-directed RNA polymerase L, 2. Fusion glycoprotein, 3. Protein V, 4. Hemagglutinin - neuraminidase, 5. Nucleocapsid (Nucleocapsid protein), 6. Phosphoprotein (Protein P), 7. Matrix protein, 8. Orf OP-2 pot. P protein, 9. Orf OP-2’, 10. (NDV) L protein, 11. Paramyxovirinae protein V zinc-binding domain-containing protein, 12. W protein. All virus proteins sequences (FASTA format) were downloaded from the UniProt (https://www.uniprot.org/), and 3D structure were generated by homology modelling using the SWISS-MODEL server (https://swissmodel.expasy.org/).^22,23^

### Mucin types in Chicken and Quail

Similar to the identification of NDV proteins, the various type of mucins reported in UniProt database were searched using keywords; Mucins, Gallus Gallus or quail, and following review for duplicates and close isoforms, sixteen types of mucins in quails and eight types of mucins in chickens were selected for analysis in this study. In case of quail mucins we looked at mucins types reported for the following breeds of quails: Coturnix japonica (CJ), Callipepla squamata (CS), Odontophorus gujanensis (OG), Coturnix pectoralis (CP).^11,12^

### Quantification of interactions between NDV proteins and chicken/quail mucins

The sequence of NDV proteins, and chicken and quail mucins were collected from UniProt database, and their 3D structures if not available as alpha fold (AF), were generated using the Swiss homology modelling (HM) tools reported before. The 3D structure of the NDV proteins, chicken and quail mucins generated in PDB format were imported onto the Chimera software and the number of hydrogen bonds (H-bond) formed between them at 10 Armstrong (10A) distance was detected. A heatmap of the number of H-bonds formed between NDV proteins and different mucins was generated to identify the high affinity interactions.

### Binding site analysis of NDV proteins, chicken, and quail mucins

The binding sites of NDV proteins and mucins of chicken and quail were identified utilizing the PrankWeb: Ligand Binding Site Prediction tool (https://prankweb.cz/). The binding sites sequences of each of NDV proteins, chickens and quail mucins were extracted from the output files and their 3D structures were generated using the Chimera software. These 3D structures of chickens and quail mucins were tested for interaction with basic 3D structures of the NDV binding sites using Chimera software and the number of hydrogen bonds (H-bond) formed is presented as heat maps.^22^

## Results

The twelve Newcastle disease virus (NDV) proteins were assessed for their interactions with all sixteen types of quail mucins by the formation of intermolecular hydrogen bonds (HB) are 10 Armstrong distance. The interaction of NDV proteins with various types of quail mucins is summarised in table 1. The number of intermolecular hydrogen bonds observed varied from 0 to 12858. The highest affinity (>10K HB) interactions were observed between mucins (Muc5A, Muc5B and Muc6) and NDV hemagglutinin-neuraminidase. This is followed by interactions (8.5K to 9.3K HB) between mucins (Muc5A, Muc5B and Muc6) and NDV fusion glycoprotein. Susd2 also showed considerable interactions (8.5K HB and 7.2K HB) with NDV hemagglutinin-neuraminidase and fusion glycoprotein respectively. In addition, Muc4 showed mid-range interactions (6 to 6.6K HB) with NDV hemagglutinin-neuraminidase and fusion glycoprotein. While the other mucin types showed weak interactions (<5K HB) with all other NDV proteins. Interestingly NDV RNA-directed RNA polymerase L, Protein V, Nucleocapsid protein, Protein P, Orf OP-2 pot. P protein, L protein and Matrix protein showed either negligible or no interactions with any of the quail mucins.

**Table 1:** Interaction of Newcastle disease virus (NDV) proteins with various types of quail mucins. Proteins Names for Newcastle disease virus (NDV): 1. RNA-directed RNA polymerase L, 2. Fusion glycoprotein, 3. Protein V, 4. Hemagglutinin - neuraminidase, 5. Nucleocapsid (Nucleocapsid protein), 6. Phosphoprotein (Protein P), 7. Matrix protein, 8. Orf OP-2 pot. P protein, 9. Orf OP-2’, 10. (NDV) L protein, 11. Paramyxovirinae protein V zinc-binding domain-containing protein, 12. W protein. The numerical values (number of intermolecular hydrogen bonds at 10 Armstrong’s) in the table indicate the intensity of the interaction between the NDV proteins and the respective mucin types. AF: Alpha fold, HM: Homology model, CJ: Coturnix japonica, CS: Callipepla squamata, OG: Odontophorus gujanensis, CP: Coturnix pectoralis, Muc: Mucin, Susd2: Sushi domain containing 2.

| No. | Mucin Types | Newcastle disease virus (NDV) Proteins |  |  |  |  |  |  |  |  |  |  |  |
| --- | --- | --- | --- | --- | --- | --- | --- | --- | --- | --- | --- | --- | --- |
|  |  | 1 | 2 | 3 | 4 | 5 | 6 | 7 | 8 | 9 | 10 | 11 | 12 |
| 1 | Muc2 Q CP HM | 0 | 4179 | 0 | 3099 | 0 | 0 | 0 | 0 | 2 | 0 | 78 | 1359 |
| 2 | Muc2 Q CS AF | 0 | 2098 | 0 | 1731 | 0 | 0 | 607 | 0 | 9 | 0 | 35 | 1634 |
| 3 | Muc2 Q OG AF | 0 | 3429 | 0 | 4682 | 0 | 0 | 0 | 0 | 0 | 0 | 0 | 259 |
| 4 | Muc4 Q CJ HM | 0 | 6066 | 0 | 5290 | 0 | 0 | 0 | 0 | 14 | 0 | 23 | 1043 |
| 5 | Muc4 Q OG AF | 0 | 6490 | 0 | 6600 | 0 | 0 | 0 | 0 | 51 | 0 | 14 | 1595 |
| 6 | Muc5A Q OG AF | 0 | 9196 | 0 | 10975 | 0 | 0 | 0 | 0 | 74 | 0 | 97 | 1388 |
| 7 | Muc5B Q OG HM | 0 | 8509 | 0 | 12858 | 0 | 0 | 0 | 0 | 344 | 0 | 47 | 2407 |
| 8 | Muc6 Q CJ HM | 0 | 9358 | 0 | 12588 | 0 | 0 | 0 | 0 | 69 | 0 | 60 | 1958 |
| 9 | Muc6 Q OG AF | 0 | 3356 | 0 | 4057 | 0 | 0 | 0 | 0 | 0 | 0 | 0 | 411 |
| 10 | Muc15 Q CJ HM | 0 | 1431 | 0 | 1792 | 0 | 0 | 0 | 0 | 2 | 0 | 0 | 165 |
| 11 | Muc15 Q OG AF | 0 | 1344 | 0 | 1708 | 0 | 0 | 0 | 0 | 1 | 0 | 0 | 194 |
| 12 | Muc16 Q OG AF | 0 | 1794 | 0 | 2244 | 0 | 0 | 0 | 0 | 0 | 0 | 0 | 127 |
| 13 | Muc24 Q OG AF | 0 | 844 | 0 | 1364 | 0 | 0 | 0 | 0 | 0 | 0 | 0 | 271 |
| 14 | MucL1 Q CS AF | 0 | 959 | 0 | 1723 | 0 | 0 | 0 | 0 | 124 | 0 | 29 | 227 |
| 15 | Susd2 Q OG AF | 0 | 7260 | 0 | 8498 | 0 | 0 | 0 | 0 | 33 | 0 | 0 | 962 |
| 16 | Muc2L Q OG AF | 0 | 3727 | 0 | 3857 | 0 | 0 | 227 | 0 | 73 | 0 | 146 | 1599 |

Similarly, the twelve NDV proteins were also assessed for interactions with the eight types of chicken mucins. The interaction of NDV proteins with various types of chicken mucins is summarised in table 2. The general pattern of interaction in chickens was similar to that observed in quails. The highest affinity (>10K HB) interactions were observed between mucins (Muc5B and Muc6) and NDV hemagglutinin-neuraminidase. While mucins (Muc5B and Muc6) also showed considerable interactions with NDV fusion glycoprotein. However, unlike quails the chicken Muc4 showed mid-range interactions (7.3K HB) with only NDV fusion glycoprotein but not hemagglutinin-neuraminidase. In addition, the chicken Muc5AC also showed mid-range interactions (5.6K HB) with NDV hemagglutinin-neuraminidase. While all other mucin types showed only weak interactions (<5K HB) with the NDV proteins.

**Table 2:**
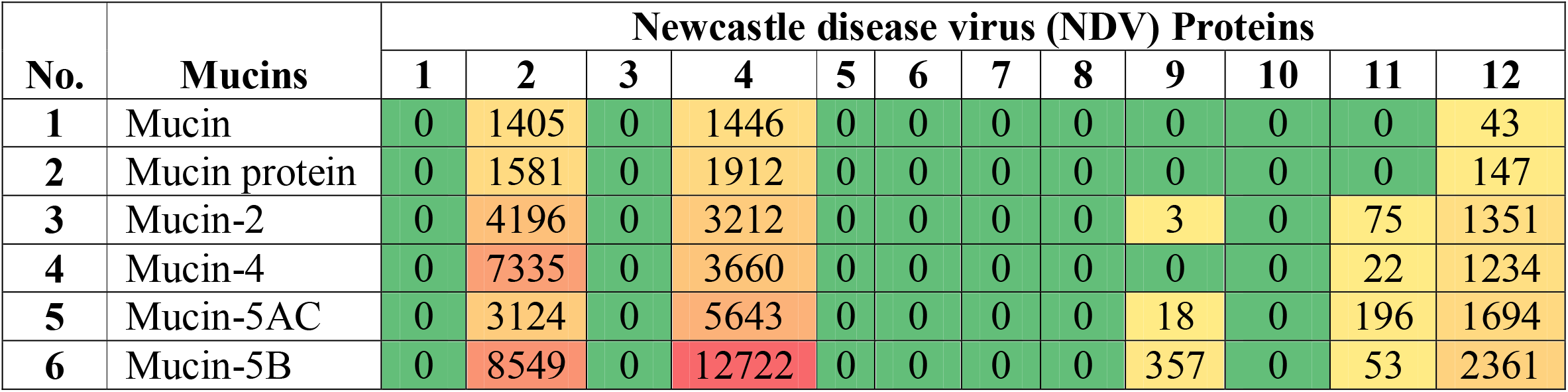
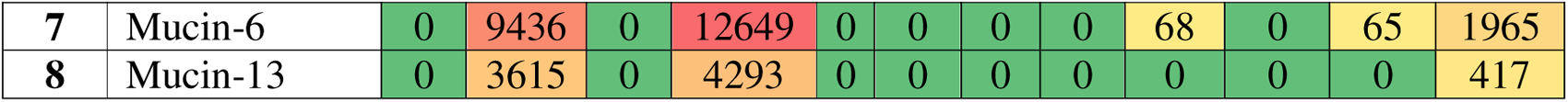
Interaction of Newcastle disease virus (NDV) proteins with various types of mucins in chicken. Proteins Names for Newcastle disease virus (NDV): 1. RNA-directed RNA polymerase L, 2. Fusion glycoprotein, 3. Protein V, 4. Hemagglutinin - neuraminidase, 5. Nucleocapsid (Nucleocapsid protein), 6. Phosphoprotein (Protein P), 7. Matrix protein, 8. Orf OP-2 pot. P protein, 9. Orf OP-2’, 10. (NDV) L protein, 11. Paramyxovirinae protein V zinc-binding domain-containing protein, 12. W protein. Mucin (UniProt ID: FIDFK7), Mucin Protein (UniProt ID: Q70YQ8).

We next assessed the number of binding sites on NDV proteins and chicken/quail mucins. The number of binding sites observed in the chicken/quail mucins was highly variable. Several of the chicken/quail mucins and NDV proteins didn’t show any binding sites and hence this was consistent with their negligible or no interactions with most NDV proteins. The chicken/quail mucins and NDV proteins which showed presence of binding sites are shown in figure 1. Among all the mucin types in chickens Muc5B showed the highest number (20) of binding site, which was followed by 13 binding sites each in Muc4 and Muc 6 (figure 1). In the quails, we observed considerable variations in the number of binding sites. While the quail Muc5B and Muc6 showed 51 and 34 binding sites respectively, the Muc5A and Susd2 had only 19 and 7 binding sites respectively (figure 1). Among the 12 NDV proteins analysed only fusion glycoprotein and hemagglutinin-neuraminidase showed presence of binding sites with 65 and 41 binding sites respectively (figure 1).

**Figure 1:**
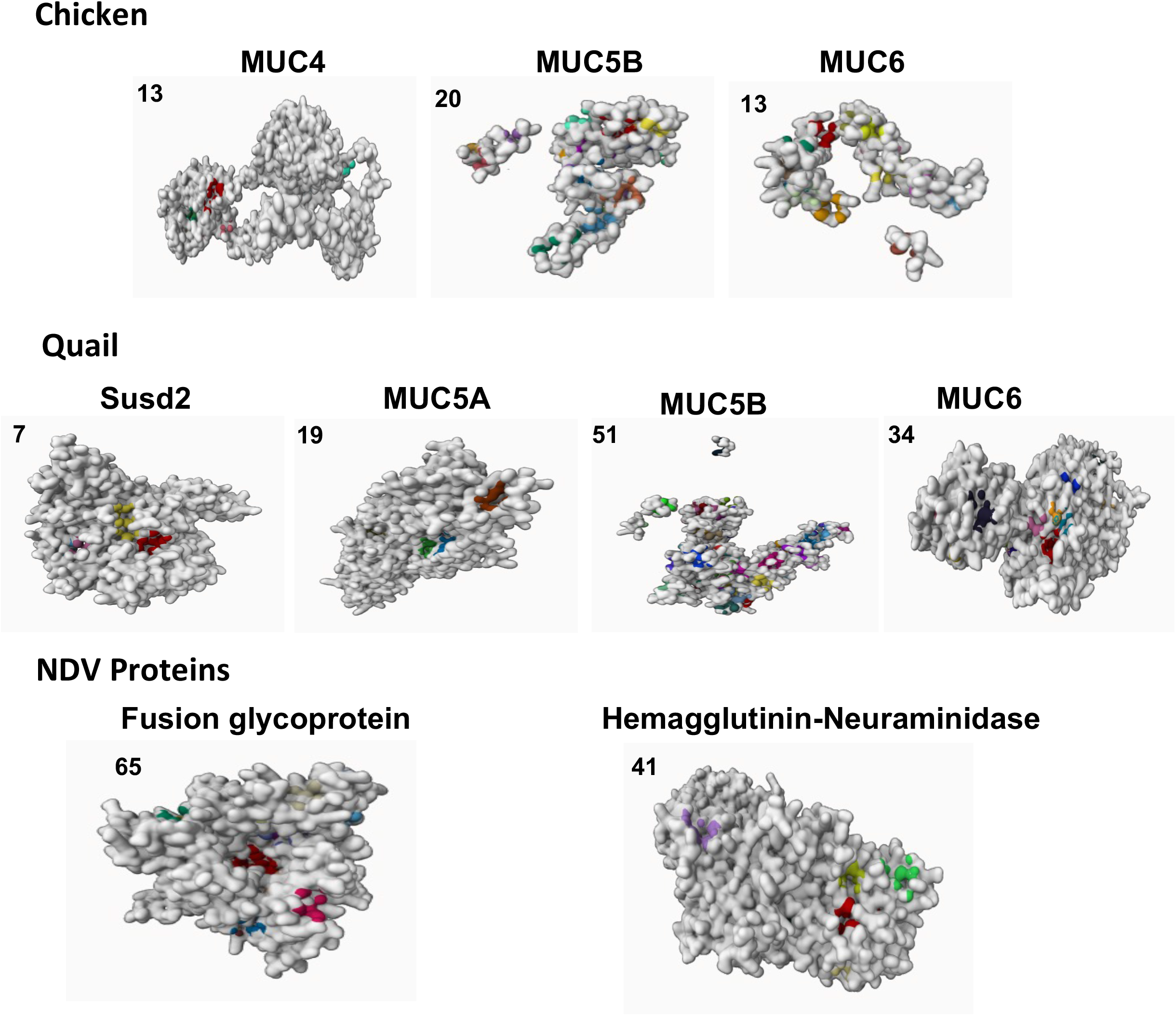
Binding sites of chicken/quail mucins and Newcastle disease virus (NDV) proteins. Representative images showing the binding sites (coloured spots) of chicken mucins (Muc4, Muc5B and Muc6), quail mucins (Susd2, Muc5A, Muc5B and Muc6) and NDV proteins (Fusion glycoprotein and Hemagglutinin-neuraminidase). The number on the top left corner of each image indicates the number of binding sites observed.

The binding sites of NDV proteins and chicken/quail mucins with the highest score were selected to further assess their interactions. While the mucins and hemagglutinin-neuraminidase showed one major binding sites each, the NDV fusion glycoprotein showed two binding sites with similar highest scores, hence both these forms (fusion glycoprotein 1 and 2) of binding sites were analysed for interactions with the mucins. The amino acid sequences of all the binding sites identified are summarised in table 3. The interactions of analysis of the major binding site of NDV proteins with chicken/quail mucins is summarised in figure 2. Muc5B was observed to have major affinity with NDV hemagglutinin-neuraminidase in chickens, while both Muc5B and Muc6 was observed to have high affinity with NDV hemagglutinin-neuraminidase in quails (figure 2). The relatively least affinity was observed between fusion glycoprotein 1 and Muc 6 in chickens and fusion glycoprotein 2 and Susd2 in quails (figure 2).

**Table 3:**
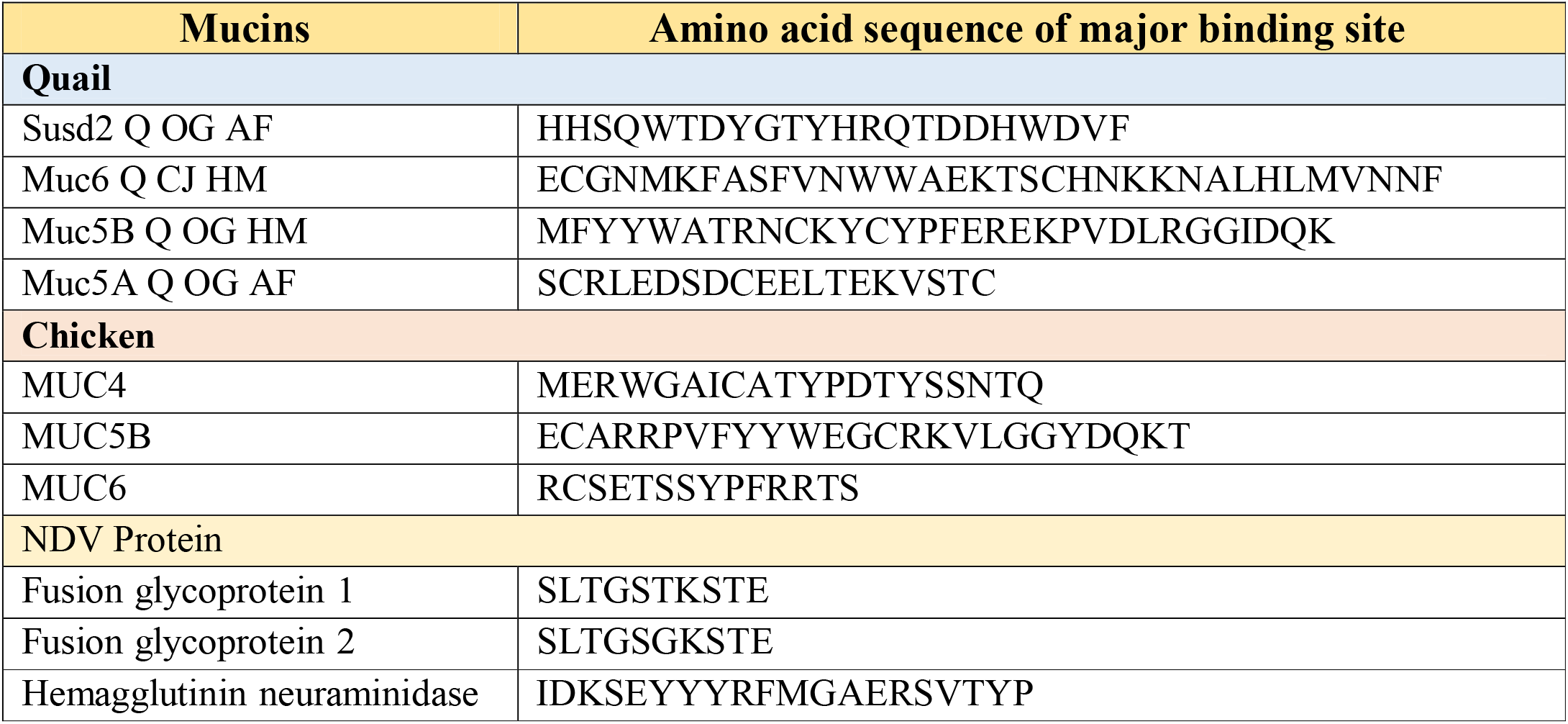
Binding sequences of Quail / Chicken Mucins and NDV Proteins.

**Figure 2:**
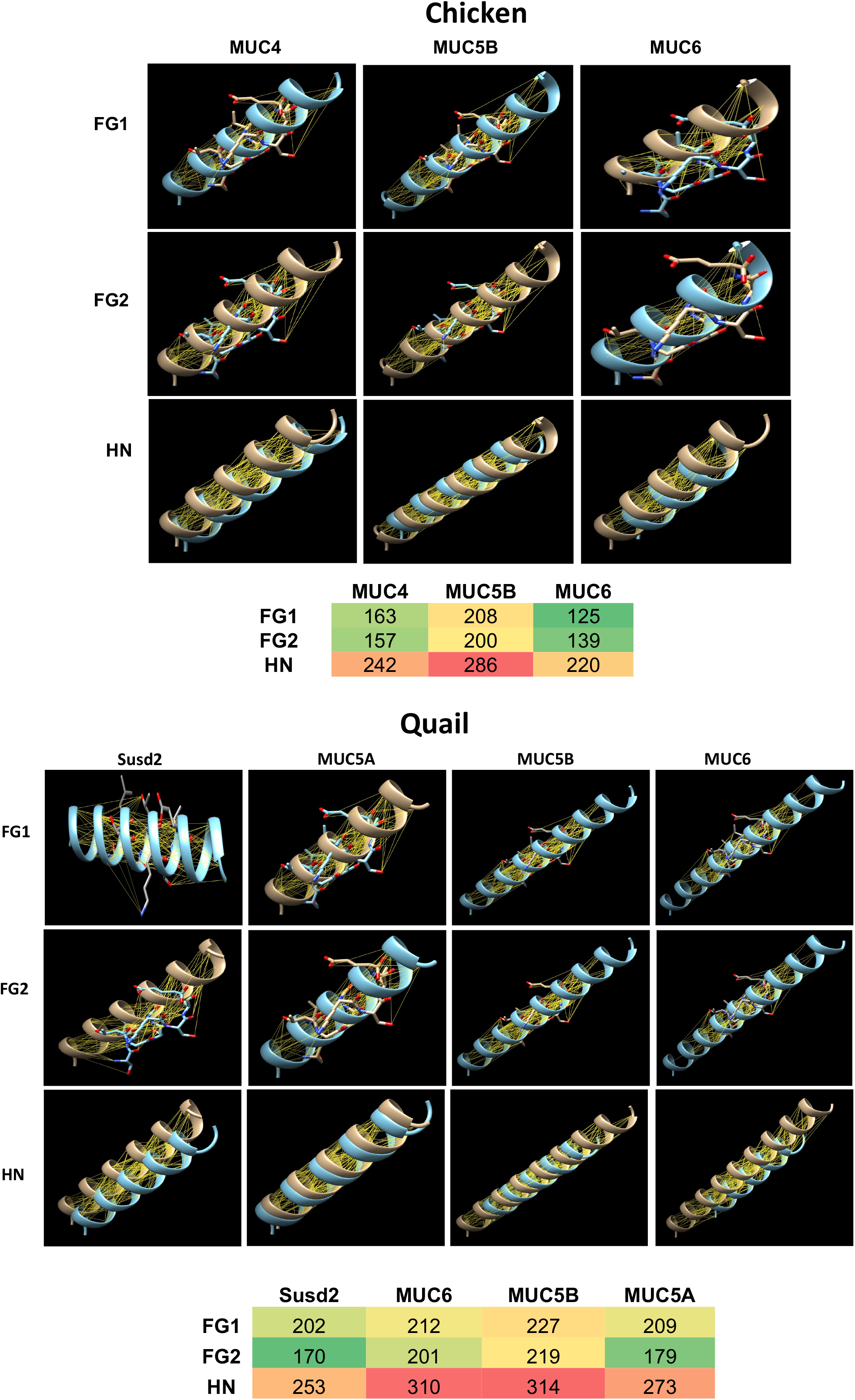
Interactions between binding sites of chicken/quail mucins with Newcastle disease virus (NDV) proteins. Representative images of interactions (hydrogen bonds, yellow lines) between chicken/quail mucins with Newcastle disease virus (NDV) proteins (Fusion glycoprotein 1 (FG1) / 2 (FG2) and Hemagglutinin-neuraminidase) is shown. The heat maps shown the number of hydrogen bonds formed between each interaction pair evaluated at 10 Armstrong distance.

## Discussion

The study provides a comprehensive analysis of the interactions between twelve Newcastle disease virus (NDV) proteins and sixteen quail and eight chicken mucins. The assessment involved the identification of intermolecular hydrogen bonds (HB) within a 10 Armstrong distance, between NDV proteins and mucins including assessment of their major binding site sequences. The findings reveal intriguing patterns of affinity and variability in binding sites, which has potential implications for developing targeted prophylaxis and therapeutics approaches based on selective targeting of the molecular interactions between NDV and host mucins in quails and chickens.

The highest affinity interactions (>10K HB) observed between NDV hemagglutinin-neuraminidase and specific quail mucins (Muc5A, Muc5B, and Muc6) suggests a strong association between these mucins and the viral entry and exit processes. This is largely consistent with extensive literature on this topic.^13,14,24-27^ Considerable interactions (8.5K to 9.3K HB) were also observed between quail mucins (Muc5A, Muc5B, and Muc6) and NDV fusion glycoprotein. This reinforces the importance of these mucins in the fusion process during viral entry, which to best of our knowledge is not previously reported. The new host-pathogen interaction insight gained from this study will prove to be useful in development of effective prevention strategies targeting mucins, which remains to be validated through preclinical studies. The general pattern of interactions in chickens mirrored the general features observed in quails, indicating consistency in the affinity of specific mucins with NDV proteins. Muc5B and Muc6 showed the highest affinity with NDV hemagglutinin-neuraminidase. This similarity feature of host-pathogen interaction is not previously reported and in our opinion merits preclinical evaluation for development of effective cross-species prevention strategies. Such cross-species prevention strategies will prove to be financially and practically viable in clinical veterinary practice and should be a focus of one health research.

Unlike quails, chicken Muc4 exhibited mid-range interactions (7.3K HB) only with NDV fusion glycoprotein, suggesting potential differences in the viral entry mechanisms between quails and chickens. While the specific biological implications of this discrepancy remain unclear,^28-30^ further investigation and validation of this observation using real-world clinical data could provide valuable insights for optimizing diagnostic and therapeutic approaches for Newcastle disease virus (NDV) infections. Understanding the unique interaction pattern of chicken Muc4 with NDV fusion glycoprotein could aid in developing specific diagnostic markers.^30,31^ This information may contribute to differentiating NDV infections in chickens from other avian diseases with similar clinical presentations. The identified interaction could also serve as a basis for developing targeted diagnostic assays. By focusing on the interaction between chicken Muc4 and NDV fusion glycoprotein, diagnostic tools can be designed to detect specific biomarkers associated with this interaction, enhancing the accuracy and specificity of NDV diagnosis in chickens.^30,31^ Investigating the precise mechanisms underlying the interaction between chicken Muc4 and NDV fusion glycoprotein may reveal vulnerabilities in the viral entry process that can be exploited for therapeutic development. To strengthen the relevance of these findings, real-world clinical data should be gathered and analysed. Comparative studies involving clinical cases of NDV infections in quails and chickens can provide valuable information on the consistency and significance of the observed mucin-virus interactions in this study. Large-scale epidemiological studies can also help establish correlations between the identified mucin-virus interactions and the prevalence, severity, and outcomes of NDV infections in different avian populations. In addition, it will be helpful to assess the tissue specific protein expression pattern of mucins in quails and chickens^11,12^ and correlate it with NDV infection pattern. Comparative expression profiling can identify mucins that are preferentially expressed in tissues targeted by NDV, providing valuable information for understanding the infection patterns. Correlating mucin expression patterns with NDV infection patterns can reveal potential host factors influencing susceptibility or resistance. This information can guide the development of interventions targeting specific tissues or mucins crucial for the virus’s successful entry and replication.

The number of binding sites on mucins and NDV proteins was highly variable, underscoring the complexity of the host-virus interactions. Muc5B in chickens stands out with the highest number (20) of binding sites, emphasizing its potential significance in the infection process. Some mucins and NDV proteins showed no binding sites, aligning with their negligible or no interactions. This indicates a degree of specificity in the interaction between certain mucins and NDV proteins.^8,17,32,33^ The major binding sites, selected based on the highest scores, reveal distinct affinities between specific mucins and NDV proteins. Muc5B exhibited major affinity with NDV hemagglutinin-neuraminidase in chickens, while Muc5B and Muc6 showed high affinity with NDV hemagglutinin-neuraminidase in quails. This finding holds promising implications for vaccine development,^32,34^ as the specific interactions between Muc5B/Muc6 and NDV reported in this study can help in the design of effective vaccines with enhanced specificity and efficacy. In contrast fusion glycoprotein showed two binding sites with similar highest scores, suggesting potential different modes of interaction with mucins. Notably, the least affinity was observed between fusion glycoprotein 1 and Muc6 in chickens and fusion glycoprotein 2 and Susd2 in quails. These results suggest that focusing on Muc5B/Muc6 - hemagglutinin-neuraminidase interactions may be a viable approach for developing effective prevention or therapeutic strategies rather than targeting the fusion glycoproteins.

The observed differences in interactions between quail and chicken mucins and NDV proteins highlight potential host-specific variations in the viral infection process.^8,31,32^ Understanding these distinctions could contribute to the development of targeted therapeutics or vaccines. The identification of specific mucins with high affinity for key NDV proteins, especially hemagglutinin-neuraminidase, suggests its functional significance in viral entry, fusion, and release processes. The identified species-specific binding sites provide potential targets for therapeutic interventions, and the amino acid sequences of these sites reported in this study offer insights into the molecular details of these interactions. In conclusion, this detailed analysis provides a valuable foundation for further investigations into the molecular mechanisms underlying NDV-mucin (Muc5B/Muc6) interactions. The insights gained from this study could have implications for the development of targeted antiviral strategies and vaccine design tailored to the specific interactions between NDV proteins and host mucins in quails and chickens.

## References

1. Seal BS, King DJ, Sellers HS. The avian response to Newcastle disease virus. Developmental & Comparative Immunology. 2000;24:257–268.

2. Alexander DJ, Senne D. Newcastle disease. Diseases of poultry. 2003;11:64–87.

3. Sun J, Han Z, Qi T, Zhao R, Liu S. Chicken galectin-1B inhibits Newcastle disease virus adsorption and replication through binding to hemagglutinin–neuraminidase (HN) glycoprotein. Journal of Biological Chemistry. 2017;292:20141–20161.

4. Bossart KN, Fusco DL, Broder CC. Paramyxovirus entry. Viral Entry into Host Cells. 2013:95–127.

5. Turan N, Ozsemir C, Yilmaz A, Cizmecigil UY, Aydin O, Bamac OE, Gurel A, Kutukcu A, Ozsemir K, Tali HE. Identification of Newcastle disease virus subgenotype VII. 2 in wild birds in Turkey. BMC Veterinary Research. 2020;16:1–8.

6. Sharif A, Ahmad T, Umer M, Rehman A, Hussain Z. Prevention and control of Newcastle disease. International Journal of Agriculture Innovations and Research. 2014;3:454–460.

7. Awan MA, Otte M, James A. The epidemiology of Newcastle disease in rural poultry: a review. Avian pathology. 1994;23:405–423.

8. Getabalew M, Alemneh T, Akeberegn D, Getahun D, Zewdie D. epidemiology, Diagnosis & Prevention of Newcastle disease in poultry. Am J Biomed Sci Res. 2019;16:50–59.

9. Bulbule N, Madale D, Meshram C, Pardeshi R, Chawak M. Virulence of Newcastle disease virus and diagnostic challenges. Adv Anim Vet Sci. 2015;3:14–21.

10. Matrosovich M, Herrler G, Klenk HD. Sialic acid receptors of viruses. Sialoglyco chemistry and biology II: tools and techniques to identify and capture sialoglycans. 2015:1–28.

11. Almhanna H, AL-Mahmodi AMM, Kadhim AB, Kumar AH. Network and structural analysis of quail mucins with expression pattern of MUC1 and MUC4 in the intestines of the Iraqi Common Quail (Coturnix Coturnix). bioRxiv. 2023:2023.2007.2018.549497.

12. AL-Mamoori NAM, Almhanna H, Kadhim AB, Kilroy D, Kumar A. Finding Expression of MUC1 and MUC4 in the Respiratory System of the Iraqi Common Quail (Coturnix coturnix). bioRxiv. 2023:2023.2009. 2001.555941.

13. Hansson GC. Mucins and the microbiome. Annual review of biochemistry. 2020;89:769–793.

14. Corfield AP. Mucins: a biologically relevant glycan barrier in mucosal protection. Biochimica et Biophysica Acta (BBA)-General Subjects. 2015;1850:236–252.

15. Ecco R, Susta L, Afonso CL, Miller PJ, Brown C. Neurological lesions in chickens experimentally infected with virulent Newcastle disease virus isolates. Avian pathology. 2011;40:145–152.

16. Nallaiyan S, Abbadorai RSAJ, Sundaramoorthy S, Nelson J, Sanyasi SVV. Production and application of recombinant haemagglutinin neuraminidase of Newcastle disease virus. Asian Pacific Journal of Tropical Medicine. 2010;3:629–632.

17. Mariappan AK, Munusamy P, Kumar D, Latheef SK, Singh SD, Singh R, Dhama K. Pathological and molecular investigation of velogenic viscerotropic Newcastle disease outbreak in a vaccinated chicken flocks. Virusdisease. 2018;29:180–191.

18. Dhaygude V, Sawale G, Chawak M, Bulbule N, Moregaonkar S, Gavhane D. Molecular characterization of velogenic viscerotropic Ranikhet (Newcastle) disease virus from different outbreaks in desi chickens. Veterinary World. 2017;10:319.

19. Rahman S, Nizamani ZA, Soomro NM, Kalhoro NH, Rasool F. Velogenic viscerotropic Newcastle disease virus produces variable pathogenicity in two chicken breeds. Journal of Animal Health and Production. 2014;2:46–50.

20. Huang Z, Panda A, Elankumaran S, Govindarajan D, Rockemann DD, Samal SK. The hemagglutinin-neuraminidase protein of Newcastle disease virus determines tropism and virulence. Journal of virology. 2004;78:4176–4184.

21. Jarahian M, Watzl C, Fournier P, Arnold A, Djandji D, Zahedi S, Cerwenka A, Paschen A, Schirrmacher V, Momburg F. Activation of natural killer cells by newcastle disease virus hemagglutinin-neuraminidase. Journal of virology. 2009;83:8108–8121.

22. Kumar AH. An assessment of the human Sortilin1 protein network, its expression and targetability using small molecules. bioRxiv. 2023:2023.2004. 2013.536697.

23. Khosravi Z, Kumar AH. Analysing the role of SERPINE1 network in the pathogenesis of human glioblastoma. Journal of Cancer Research. 2023;1:1–6.

24. Li Q, Wei D, Feng F, Wang X-L, Li C, Chen Z-N, Bian H. α2, 6-linked sialic acid serves as a high-affinity receptor for cancer oncolytic virotherapy with Newcastle disease virus. Journal of Cancer Research and Clinical Oncology. 2017;143:2171–2181.

25. Johansson ME, Sjövall H, Hansson GC. The gastrointestinal mucus system in health and disease. Nature reviews Gastroenterology & hepatology. 2013;10:352–361.

26. Linden S, Sutton P, Karlsson N, Korolik V, McGuckin M. Mucins in the mucosal barrier to infection. Mucosal Immunol 1: 183–197. In; 2008.

27. Rose MC, Voynow JA. Respiratory tract mucin genes and mucin glycoproteins in health and disease. Physiological reviews. 2006;86:245–278.

28. Lewis AL, Lewis WG. Host sialoglycans and bacterial sialidases: a mucosal perspective. Cellular microbiology. 2012;14:1174–1182.

29. Dharmani P, Srivastava V, Kissoon-Singh V, Chadee K. Role of intestinal mucins in innate host defense mechanisms against pathogens. Journal of innate immunity. 2009;1:123–135.

30. Chaturvedi P, Singh AP, Batra SK. Structure, evolution, and biology of the MUC4 mucin. The FASEB journal: official publication of the Federation of American Societies for Experimental Biology. 2008;22:966.

31. Liu T, Song Y, Yang Y, Bu Y, Cheng J, Zhang G, Xue J. Hemagglutinin–Neuraminidase and fusion genes are determinants of NDV thermostability. Veterinary microbiology. 2019;228:53–60.

32. Panner Selvam MK, Kanagaraj V, Kathaperumal K, Nissly RH, Daly JM, Kuchipudi SV. Comparative transcriptome analysis of spleen of Newcastle Disease Virus (NDV) infected chicken and Japanese quail: a potential role of NF-κβ pathway activation in NDV resistance. VirusDisease. 2023;34:402–409.

33. Abd El Aziem A, Abd-Ellatieff H, Elbestawy A, Belih S, El-Hamid A, Abou-Rawash A-R. Susceptibility of Japanese quail and chickens to infection with newcastle disease virus genotype VIId. Damanhour Journal of Veterinary Sciences. 2020;3:27–31.

34. Rota P, La Rocca P, Piccoli M, Montefiori M, Cirillo F, Olsen L, Orioli M, Allevi P, Anastasia L. Potent Inhibitors against Newcastle Disease Virus Hemagglutinin-Neuraminidase. ChemMedChem. 2018;13:236–240.

